# Telomere length and *TERT* expression are associated with age in almond (*Prunus dulcis* [Mill.] D.A.Webb)

**DOI:** 10.1101/2020.09.25.294074

**Authors:** Katherine M. D’Amico-Willman, Elizabeth Anderson, Thomas M. Gradziel, Jonathan Fresnedo-Ramírez

**Author notes:** Authors contributed equally (Katherine M. D’Amico-Willman, Elizabeth Anderson).

## Abstract

While it is well known that all organisms age, our understanding of how aging occurs varies dramatically among species. The aging process in perennial plants is not well defined, yet can have implications on production and yield of valuable fruit and nut crops. Almond, a relevant nut crop, exhibits an age-related disorder known as non-infectious bud failure (BF) that affects vegetative bud development, indirectly affecting kernel-yield. This species and disorder present an opportunity to address aging in a commercially-relevant and vegetatively-propagated, perennial crop threatened by an aging-related disorder. In this study, we tested the hypothesis that telomere length and/or *TERT* expression can serve as biomarkers of aging in almond using both whole-genome sequencing data and leaf samples collected from distinct age cohorts over a two-year period. To measure telomere lengths, we employed both *in silico* and molecular approaches. We also measured expression of *TERT*, a subunit of the enzyme telomerase, which is responsible for maintaining telomere lengths. Results from this work show a marginal but significant association between both telomere length measured by monochrome multiplex quantitative PCR and *TERT* expression, and age of almond seedlings. These results suggest that as almonds age, *TERT* expression decreases and telomeres shorten. This work provides valuable information on potential biomarkers of perennial plant aging, contributing to our limited knowledge of this process. In addition, translation of this information will provide opportunities to address BF in almond breeding and nursery propagation.

## Introduction

The current concept and study of aging is centered primarily around mammals with research focused on circumventing deleterious impacts on health (Kirkwood, 2005; Sanders and Newman, 2013). However, all eukaryotic organisms exhibit signals of aging, resulting in the deterioration of key biological processes and subsequent decrease in health, performance, and fitness of individuals. Perennial plants represent a unique model to address the aging process and its impact since these species undergo cycles of dormancy and growth, and maintain the ability to reproduce for multiple years. The aging process of perennial plants is relevant due to the longevity and economic importance of perennial crops such as fruit and nut trees (Munné-Bosch, 2007; Brutovská *et al*., 2013; Thomas, 2013). Individual trees can remain productive in orchards for decades; however, aging in plants and its implications for growth and reproduction are neglected areas of research with potential consequences for production, management, conservation, and breeding.

The lack of understanding of aging in perennials is partly due to the complexity in measuring and conceptualizing age in perennial plant species since aging occurs chronologically and ontogenetically in opposite directions (Poethig, 2003). Chronologic age can be defined as the amount of time since tissue/organ formation (e.g. human skin cells replenish every few days, meaning each cell is typically a day or 2-days old), while ontogenetic age refers more to developmental time and allows for the accumulation of mutations or chromosomal alterations (e.g. 2-day old skin cells at age 6 compared to 2-day old skin cells at age 60). Agriculturally relevant perennials are often vegetatively propagated (i.e. cloned), blurring the distinction between ontogenetic and chronologic age, and tend to be grown under intensive management. The limitation in determining age in perennials creates a need to identify biomarkers in these species that enable ontogenetic age estimation.

Almond (*Prunus dulcis* [Mill.] D.A.Webb; Fig. 1) is an economically relevant, Rosaceous crop, subject to intense horticultural management to maintain maximum nut production. In California, the almond industry is estimated to contribute ∼$11 billion to the state’s GDP annually (Almond Board of California, 2019). Top-producing almond cultivars, some of which were first obtained more than 100 years ago, are produced for commercial orchards via vegetative propagation (Micke, 1996; Wickson, 1914). As orchards age (after 20-25 years), trees are replaced with “new” clones of typically the same cultivar to maintain homogeneity in quality and high levels of production (Micke, 1996).

**Figure 1.**
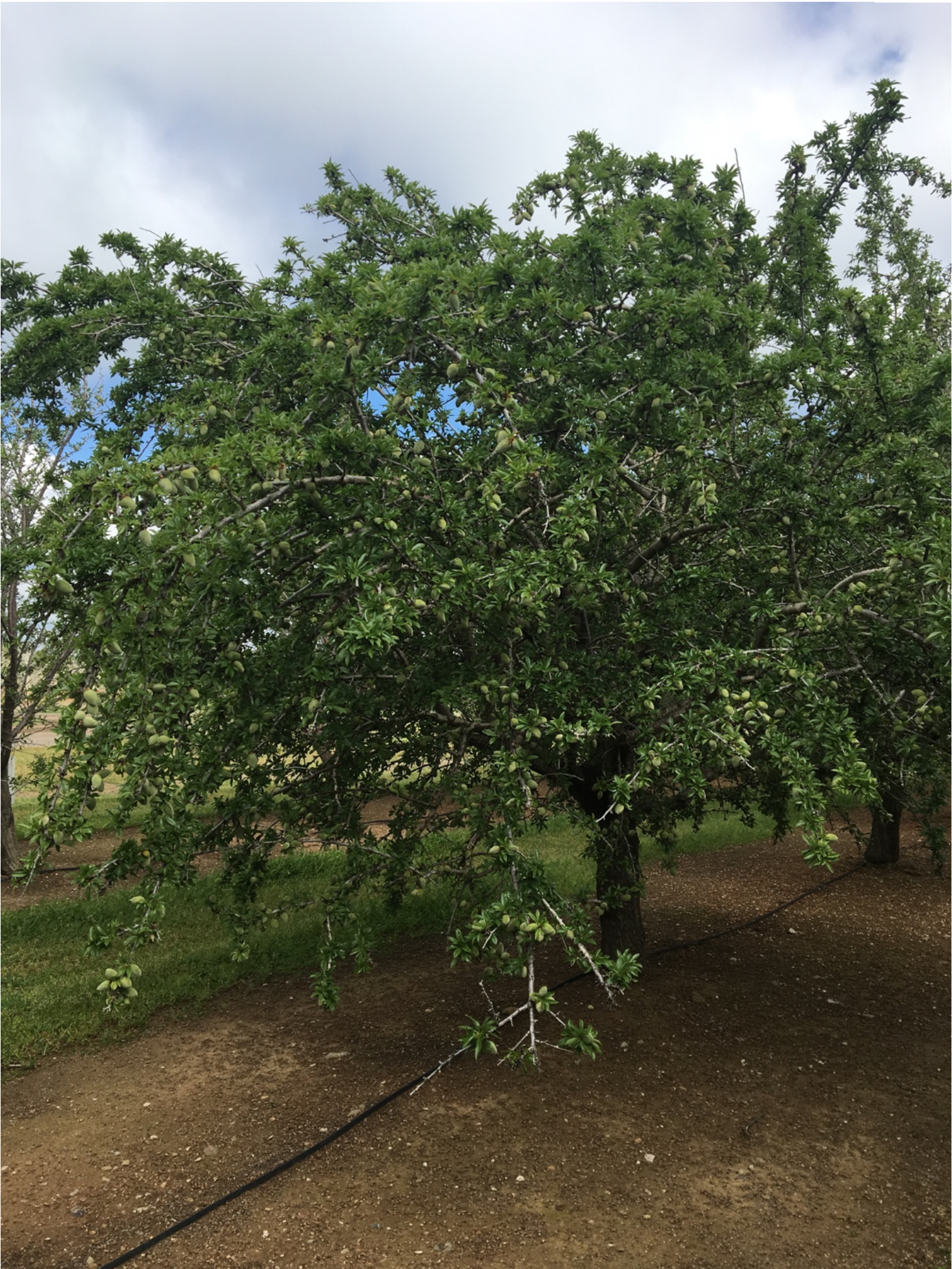
Image of almond cultivar ‘Nonpareil’ (photo taken by K. D’Amico-Willman in May 2018).

Almond exhibits an age-related disorder known as non-infectious bud-failure (BF) affecting vegetative bud development in the spring (Micke, 1996; Kester, 1970). Genotypes exhibiting this disorder show characteristic dieback at the top of the canopy, and severe levels of BF can result in up to 50% yield loss (Gradziel et al. 2013). Empirical evidence shows BF is associated with age (Kester *et al*., 2004); however, as almonds are produced primarily through vegetative propagation rather than by seed, their true ontogenetic age and thus susceptibility to BF can be difficult to assess (Micke, 1996). Biomarkers indicative of age would be valuable to growers, breeders, and producers to screen germplasm. Thus, almond represents a potential model species for the study of aging in perennials due to its economic relevance, the abundance of available germplasm and breeding programs, and the exhibition of an age-related disorder.

One biomarker of aging is telomere length measurement (Sanders and Newman, 2013; Marioni *et al*., 2016; Runov *et al*., 2015), which has been primarily studied in animals. Telomeres are nucleoproteins that cap the end of chromosomes, preventing premature instability of genomic material and cellular senescence (Watson and Riha, 2011). Telomeres tend to shorten over mitotic cellular divisions due to decreased levels of telomerase, an enzyme that supports telomere replication during the S-phase of the cell cycle (Nelson *et al*., 2014). Over mitotic cell divisions, telomeres eventually reach a critical minimum length at which point the cell senesces and dies due to genome instability resulting from single stranded DNA at the ends of chromosomes (Hemann *et al*., 2001). This progressive shortening is proposed as a marker of aging in mammalian cells and is linked to physiological deterioration and some age-related disorders (Sanders and Newman, 2013; Watson and Riha, 2011; Aviv and Shay, 2018). Plant chromosomes also contain telomeres with similar functions. While the relationship between telomeres and the aging process is not as clearly defined in plants as in animals, previous work shows associations between telomere length and various stages of plant development (Zachová *et al*., 2013; Watson and Riha, 2011; Procházková Schrumpfová *et al*., 2019) suggesting telomere length could be a suitable biomarker of age in plants.

Given that telomerase activity modulates telomere length, expression of genes involved in the telomerase biosynthetic pathway could also serve as biomarkers for aging (Fitzgerald *et al*., 1996; De la Torre-Espinosa *et al*., 2020; Boccardi and Paolisso, 2014; Anchelin *et al*., 2011; Fossel, 1998). TERT is the catalytic subunit of the telomerase enzyme (Oguchi *et al*., 1999) and the RNA subunit (TR) functions as the template for reverse transcription (Procházková Schrumpfová *et al*., 2019). Expression of *TERT* is shown to affect telomerase activity (Sweetlove and Gutierrez, 2019; Jurečková *et al*., 2017). In Arabidopsis, increased *TERT* expression is linked to proportional increases in telomerase activity and telomere length (Zangi *et al*., 2019; Fitzgerald *et al*., 1999) which is in turn linked to age. Since *TERT* is tied to telomere length in both plants and animals, its expression may also serve as an indicator of age in plants (Watson and Riha, 2011).

This study tests the hypothesis that telomere length and *TERT* expression in almond are associated with age and can thus serve as biomarkers of aging in this species. Both telomere length and *TERT* expression show promise as diagnostic biomarkers since they can be measured in a high-throughput manner via *in silico* and molecular approaches (Nersisyan and Arakelyan, 2015; Montpetit *et al*., 2014; Cawthon, 2009). These approaches build on previous research examining the relationship between telomere lengths and age in perennial plants (Flanary and Kletetschka, 2005; Moriguchi *et al*., 2007; Liang *et al*., 2015; Liu *et al*., 2007). The goal of this work is to advance our understanding and provide a model for the study of aging and its implications in perennial plant species.

## Materials and Methods

### Whole-genome Sequencing Data

To perform *in silico* mean telomere length estimation, fastq files produced from whole-genome sequencing of nine almond accessions were downloaded to the Ohio Supercomputer Center (Ohio Supercomputer Center, 1987) from the National Center for Biotechnology Information Sequence Read Archive from bioprojects PRJNA339570 and PRJNA339144. Accessions were selected for use in this study if the age of the individual was included in the metadata for the biosample entry on NCBI SRA. Table 1 includes the SRA biosample number for each almond accession used as well as accession name, cultivar name (if available), and age.

**Table 1.** NCBI SRA biosample entry for almond accessions used for *in silico* mean telomere length estimation.

| SRA Biosample | Accession Name | Cultivar | Age (years) |
| --- | --- | --- | --- |
| SRR4045222 | DPRU 1207.2 | N/A | 29 |
| SRR4045223 | DPRU 210 | ‘Languedoc’ | 32 |
| SRR4045224 | DPRU 2331.9 | N/A | 20 |
| SRR4045225 | DPRU 1791.3 | N/A | 23 |
| SRR4045226 | DPRU 2301 | ‘Tuono’ | 21 |
| SRR4045227 | DPRU 2374.12 | N/A | 19 |
| SRR4045228 | DPRU 1456.4 | ‘Badam’ | 28 |
| SRR4045229 | DPRU 1462.2 | N/A | 28 |
| SRR4036105 | DPRU 2578.2 | N/A | 15 |

### In silico Mean Telomere Length Estimation

Mean telomere lengths were estimated for nine almond accessions using files in fastq format containing whole-genome sequencing data for each individual. These fastq files were used as input in the program Computel v. 1.2 (Nersisyan and Arakelyan, 2015). To estimate mean telomere lengths, the computel.sh script was run with the following parameters: -nchr 8 - lgenome 227411381 -pattern CCCTAAA -minseed 14 using the estimated length of the peach genome based on the v2 assembly (Initiative *et al*., 2013). All work was performed using the Ohio Supercomputer Center computational resources (Ohio Supercomputer Center, 1987).

### Plant Material

Leaf samples for this study were collected in May 2018 and 2019 from almond breeding selections located at the Wolfskill Experimental Orchards (Almond Breeding Program, University of California – Davis, Winters, CA). Tissue was harvested from the canopy of a total of 36 unique individuals representing distinct age cohorts (Table 2). Samples were immediately frozen on ice and stored at −20 °C until shipment overnight on dry ice to the Ohio Agricultural Research and Development Center (OARDC – Wooster, OH). Samples were stored at −20 °C until processing, and all subsequent experimental procedures were conducted at the OARDC.

**Table 2.** Sampling scheme for 2018 and 2019 almond age cohort collections.

| Sampling Year | Age (years) | Sample Size |
| --- | --- | --- |
| 2018 | 1 | 6 |
| 2018 | 5 | 6 |
| 2018 | 9 | 6 |
| 2018 | 14 | 6 |
| 2019 | 2 | 4 |
| 2019 | 7 | 4 |
| 2019 | 11 | 4 |

### DNA and RNA Extraction

DNA was extracted from the age cohort samples using the Omega E-Z 96® Plant DNA Kit (Omega Bio-tek, Norcross, GA) with slight modification. Briefly, 100 mg of leaf material was weighed in 2.0 mL tubes containing two 1.6 mm steel beads and kept frozen in liquid nitrogen. Samples were ground in a 2000 Geno/Grinder*®* (SPEX SamplePrep, Metuchen, NJ) in two 48-well cryo-blocks frozen in liquid nitrogen. Following a 65 °C incubation, samples were incubated on ice for 20 minutes, treated with 10 μL of RNase solution (2.5 ul RNase [Omega Bio-tek, Norcross, GA] + 7.5 μl TE pH 8), equilibrated through addition of 150 μl Equilibration Buffer (3 M NaOH), incubated at room temperature for four minutes, and centrifuged at 4,400 rpm for two minutes prior to the addition of SP3 buffer. Concentration and quality were analyzed using a NanoDrop*™* 1000 spectrophotometer and a Qubit 4 Fluorometer with a dsDNA HS Assay Kit (ThermoFisher Scientific, Waltham, MA).

RNA was extracted following the protocol outline in Gambino *et al*. (2008) with slight modifications. Briefly, leaf material was ground in liquid nitrogen using a mortar and pestle, and 150 mg of tissue was weighed into a 2.0 mL microfuge tube frozen in liquid nitrogen. To extract RNA, 900 μL CTAB extraction buffer (2% CTAB, 2.5% PVP-40, 2 M NaCl, 100 mM Trish-HCl pH 8.0, 25 mM EDTA pH 8.0, 2% Beta-mercaptoethanol added before use) was added to each tube and samples were incubated at 65 °C for ten minutes. Following incubation, two phase separations were performed using an equal volume of chloroform:isoamyl alcohol (24:1). RNA was precipitated in 3 M lithium chloride and incubated on ice for 30 minutes, and samples were pelleted by centrifugation at 21,000 x g for 15 minutes. Pellets were then resuspended in 500 μL pre-warmed SSTE buffer (10 mM Tris-HCl pH 8.0, 1 mM EDTA pH 8.0, 1% SDS, 1 M NaCl) followed by a phase separation with an equal volume of chloroform:isoamyl alcohol (24:1). A final precipitation was performed using 0.7 volumes chilled 100% isopropanol. RNA was pelleted and washed with 70% ethanol before being resuspended in 30 μL nuclease-free water. A DNase treatment was performed using DNA-*free™* DNA Removal Kit (ThermoFisher Scientific) according to the manufacturer’s instructions. All materials used for extraction were nuclease-free and cleaned with RNaseZap*™* RNase decontamination wipes (ThermoFisher Scientific) prior to use. All centrifugation steps were performed at 4°C. RNA quality and concentration were assessed using a NanoDrop*™* 1000 spectrophotometer and a Qubit 4 Fluorometer with an RNA HS Assay Kit (ThermoFisher Scientific).

### Monochrome Multiplex Quantitative PCR (MMQPCR) to Measure Relative Telomere Lengths

MMQPCR was conducted following the protocol outlined in Vaquero-Sedas and Vega-Palas (2014) with minimal modifications. Primer sequences for genes used in this study are shown in Table 3, including primers for the single copy gene, *PP2A*, and for the telomere sequence (Wang *et al*., 2014; Vaquero-Sedas and Vega-Palas, 2014). Oligos were synthesized by MilliporeSigma (Burlington, MA) and resuspended to a concentration of 100 μM upon arrival. Standard curves were created for each primer pair by pooling six aliquots of DNA isolated from a single clone of the almond cultivar ‘Nonpareil’, and performing successive dilutions to 20 ng/μL, 10 ng/μL, 1 ng/μL, 0.5 ng/μL, and 0.25 ng/μL. Reactions were carried out in triplicate for each primer by concentration combination.

**Table 3.**
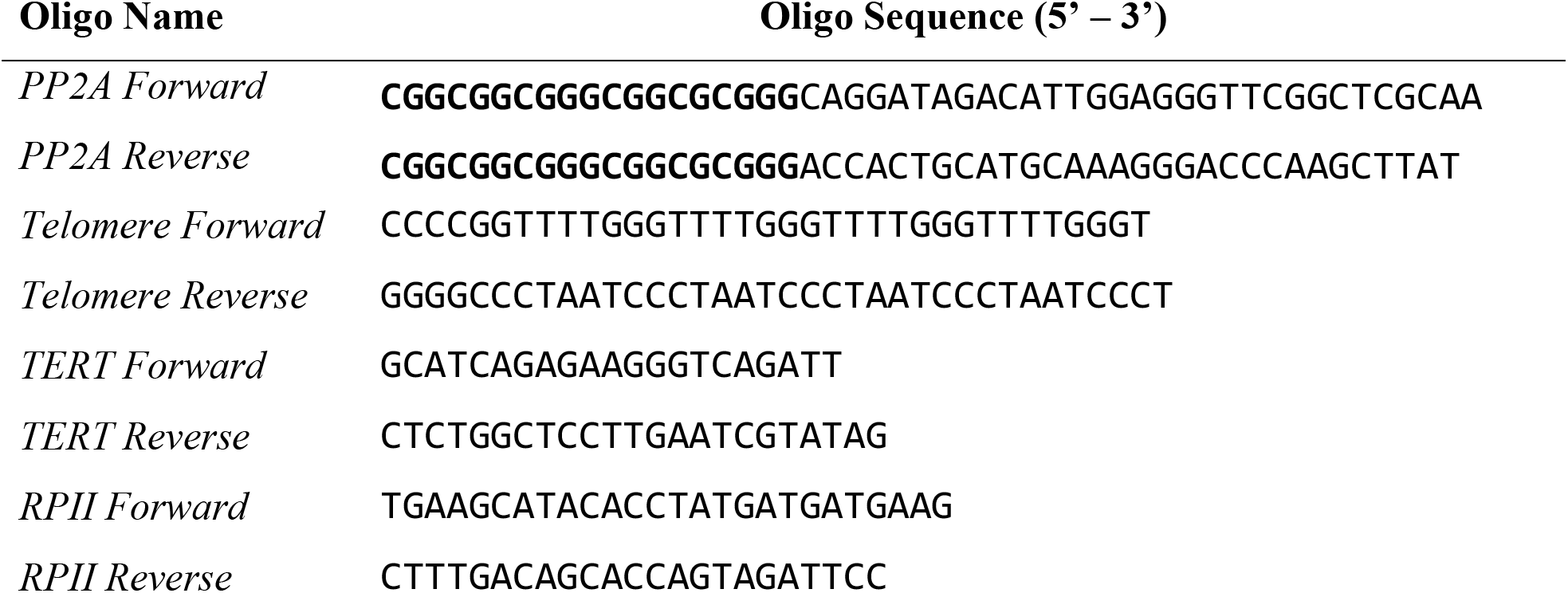
Oligos used for all MMQPCR and qRT-PCR studies.

Isolated DNA from the age cohort samples was diluted to 20 ng/μL. Multiplex reactions were carried out in sextuplicate for each replicate within the age cohorts in a 10 μL volume using QuantaBio PerfeCTa SYBR® Green SuperMix (Quanta Biosciences, Beverly, MA) (2X), forward and reverse primers (100 nM each), and 20 ng template DNA according to the manufacturer’s instructions. Reactions were performed in a Bio Rad C1000 Touch Thermal Cycler (Bio Rad Laboratories, Hercules, CA) using the following program: initial denaturation at 95 °C for 3 minutes followed by 2 cycles of incubation at 94 °C for 15 seconds and annealing at 49 °C for 15 seconds; telomere and *PP2A* amplicons were generated following 35 cycles at 95 °C for 30 seconds, 59 °C for 1 minute, 72 °C for 30 seconds, 84 °C for 15 seconds and 85 °C for 15 seconds; final incubation at 72 °C for 1 minute. Melting curve analysis was performed at a temperature range of 74-85 °C for both primer pairs to ensure no non-specific amplification.

### cDNA Synthesis and Quantitative Reverse Transcriptase PCR (qRT-PCR) to Measure Relative Expression of TERT

Reactions were carried out in a 20 μL volume using the Verso™ cDNA synthesis Kit (ThermoFisher Scientific). One reaction was prepared for each age cohort sample according to the manufacturer’s instructions. Reactions were performed in a MJ Research PTC-200 thermal cycler using the following program: 42 °C for 30 minutes followed by 95 °C for 2 minutes.

To quantify expression of *TERT* in age cohort individuals, qRT-PCR was performed in triplicate for each sample. The gene *RPII* from peach was used as a reference (Bastias *et al*., 2020; Tong *et al*., 2009), and the sequence for the *TERT* gene was derived from the ‘Texas’ genome (https://www.rosaceae.org/analysis/295) using the homologous peach gene sequence as a reference (Alioto *et al*., 2020). Primer sequences are shown in Table 2, and all oligos were synthesized by MilliporeSigma (Burlington, MA) and resuspended to a concentration of 100 μM upon arrival.

To generate cDNA from the age cohort samples, 100 ng of RNA was used as input in the Verso cDNA Synthesis Kit (ThermoFisher Scientific) according to the manufacturer’s instructions. To test for relative expression of *TERT*, reactions were carried out in triplicate for each biological replicate within the age cohorts in a 10 μL volume using QuantaBio PerfeCTa SYBR® Green SuperMix (Quanta Biosciences) (1X), forward and reverse primers (100 nM), and cDNA (1 μL) according to the manufacturer’s instructions. Reactions were performed in Bio Rad C1000 Touch Thermal Cycler (Bio Rad Laboratories) using the following program: initial denaturation at 95 °C for 3 minutes followed by 40 cycles at 95 °C for 15 seconds and 55 °C for 45 seconds. Melt curves were generated at a temperature range of 74-85 °C for both primer pairs to ensure no non-specific amplification.

### Statistical Analysis

Mean telomere lengths generated *in silico* using Computel v. 1.2 for each almond accession were square root transformed prior to analysis. Mean telomere length was regressed on chronological age to test for a linear relationship based on *in silico* predictions. Normality was confirmed using a Shapiro-Wilks test. Mean telomere lengths generated for each individual almond accession are listed in Supplementary File S1.

Using the standard curve generated with *PP2A* (S) and telomere (T) primers for a reference almond sample, relative T/S ratios were calculated for each individual sample based on Cq values for the telomere and *PP2A* products (Vaquero-Sedas and Vega-Palas, 2014). Z-scores were calculated from the T/S ratios as recommend in Verhulst (2020) for each replicate within the age cohorts. Normality and homogeneity of variance were confirmed using Shapiro-Wilks and Bartlett tests. Analysis of variance (ANOVA) was performed for each age cohort followed by *post hoc* Fisher’s LSD and pairwise t-tests. Gene expression data were analyzed according to guidelines in Bustin et. al (2009), first by normalizing *TERT* expression to that of the reference gene, *RPII*. Following normalization, data were log-transformed, and normality and homogeneity of variance were confirmed using Shapiro-Wilks and Bartlett tests. ANOVA was performed for each age cohort followed by *post hoc* analysis with Tukey’s HSD. All analyses were performed using R v. 3.6.1 and plots were generated using ggplot2 v. 3.3.0. Calculated T/S ratios, relative telomere lengths, relative *TERT* expression and log-transformed *TERT* expression as well as raw Cq values for each individual are listed in Supplementary File S1. All R code used to perform analyses is reported in Supplementary File S2.

## Results

### Associations of telomere length and age in almond

Mean telomere lengths were estimated *in silico* using whole-genome sequencing data for nine select almond accessions and regressed against accession age following a square root transformation. Normality of residuals was confirmed using a Shapiro-Wilks test (p-value = 0.318). Linear regression suggests a negative relationship between mean telomere length and age as depicted in Fig. 2.

**Figure 2.**
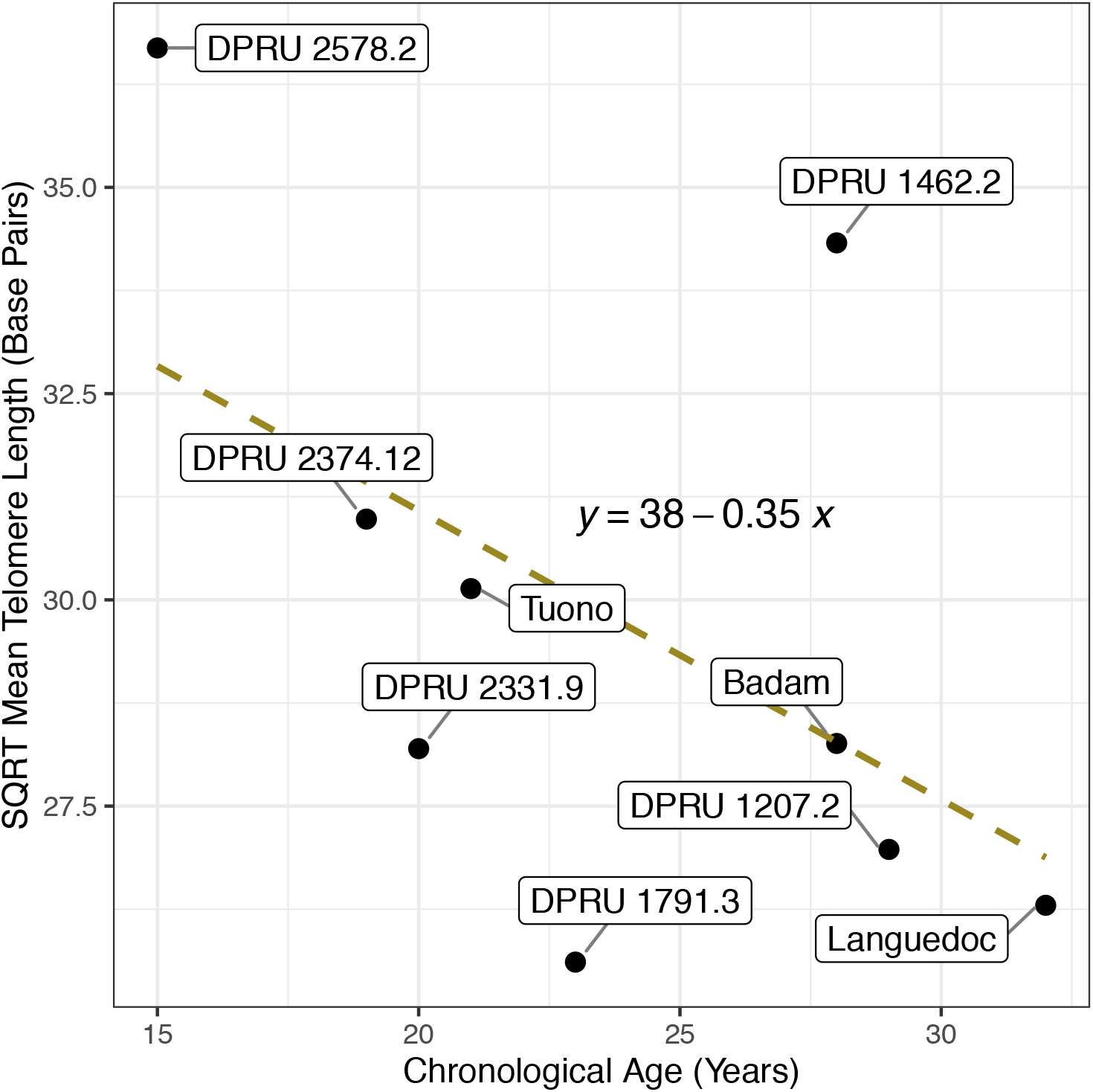
Linear regression showing the relationship between 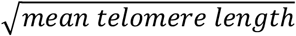 and accession age in nine almond accessions. Dashed line represents the best-fit linear model (p-value = 0.1458; R^2^ = 0.2767).

Relative telomere lengths were generated for the almond individuals within each of the age cohorts collected in 2018 (1, 5, 9, and 14 years) and 2019 (2, 7, and 11 years old) using the MMQPCR approach. Normality of residuals and homogeneity of variance of relative telomere lengths were confirmed using Shapiro-Wilks (**2018:** p-value = 0.2578, n = 4-6; **2019:** p-value = 0.4682, n = 3) and Bartlett (**2018:** p-value = 0.1408; **2019:** p-value = 0.4613) tests. ANOVA results for the linear model, z-score ∼ age, were marginally significant in both 2018 and 2019, and subsequent *post hoc* Fisher’s LSD and pairwise t-tests revealed significant differences between ages 1 and 14 years and 5 and 14 years (Fig. 3a) in the 2018 cohorts, and between ages 2 and 11 years old (Fig. 3b) in the 2019 cohorts.

**Figure 3.**
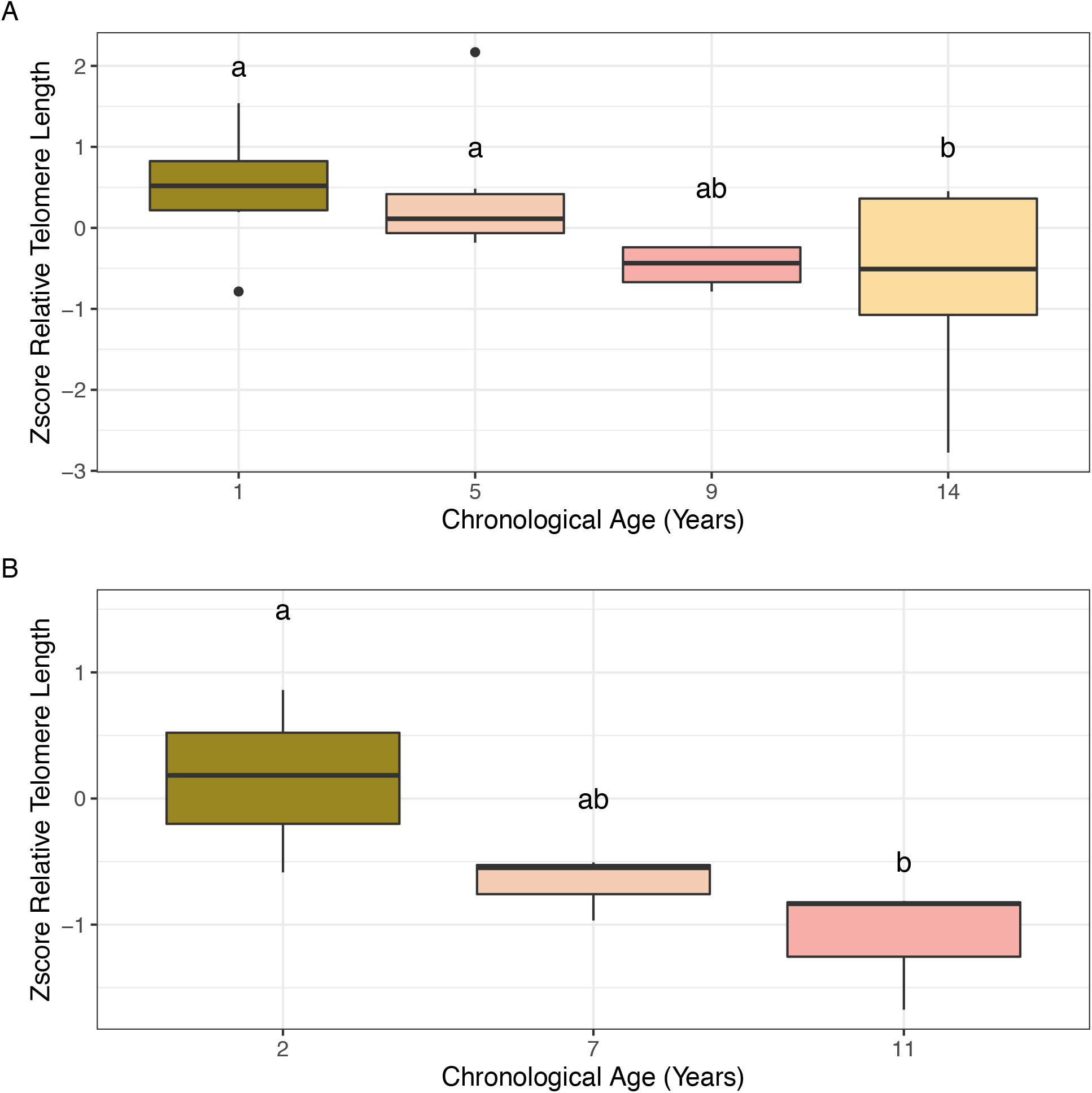
Boxplots depicting the calculated z-score of the T/S ratio for almond samples within the age cohorts tested. **(a)** Age cohort collected in 2018. **(b)** Age cohort collected in 2019. Significant differences in z-scores between age cohorts based on ANOVA followed by *post hoc* Fisher’s LSD (*α* = 0.1) are denoted by letter groupings (ANOVA 2018 p-value = 0.1077; ANOVA 2019 p-value = 0.06548).

### TERT gene expression patterns associated with age in almond

Normalized expression of *TERT* was measured for almond samples among the age cohorts collected in 2018 and 2019 for this study using *RPII* as the reference gene. Normality of residuals and homogeneity of variance were confirmed using Shapiro-Wilks (**2018:** p-value = 0.694, n = 2-3; **2019:** p-value = 0.09456, n = 4) and Bartlett (**2018:** p-value = 0.6976; **2019:** p-value = 0.3579) tests. ANOVA results comparing the average log(expression) values for each age cohort revealed significant differences between cohorts in both 2018 and 2019. *Post hoc* analysis with Tukey’s HSD revealed significant differences in *TERT* expression between ages 1 and 14 years old in the 2018 cohorts (Fig. 4a) and between ages 2 and 11 years old in the 2019 age cohorts (Fig. 4b).

**Figure 4.**
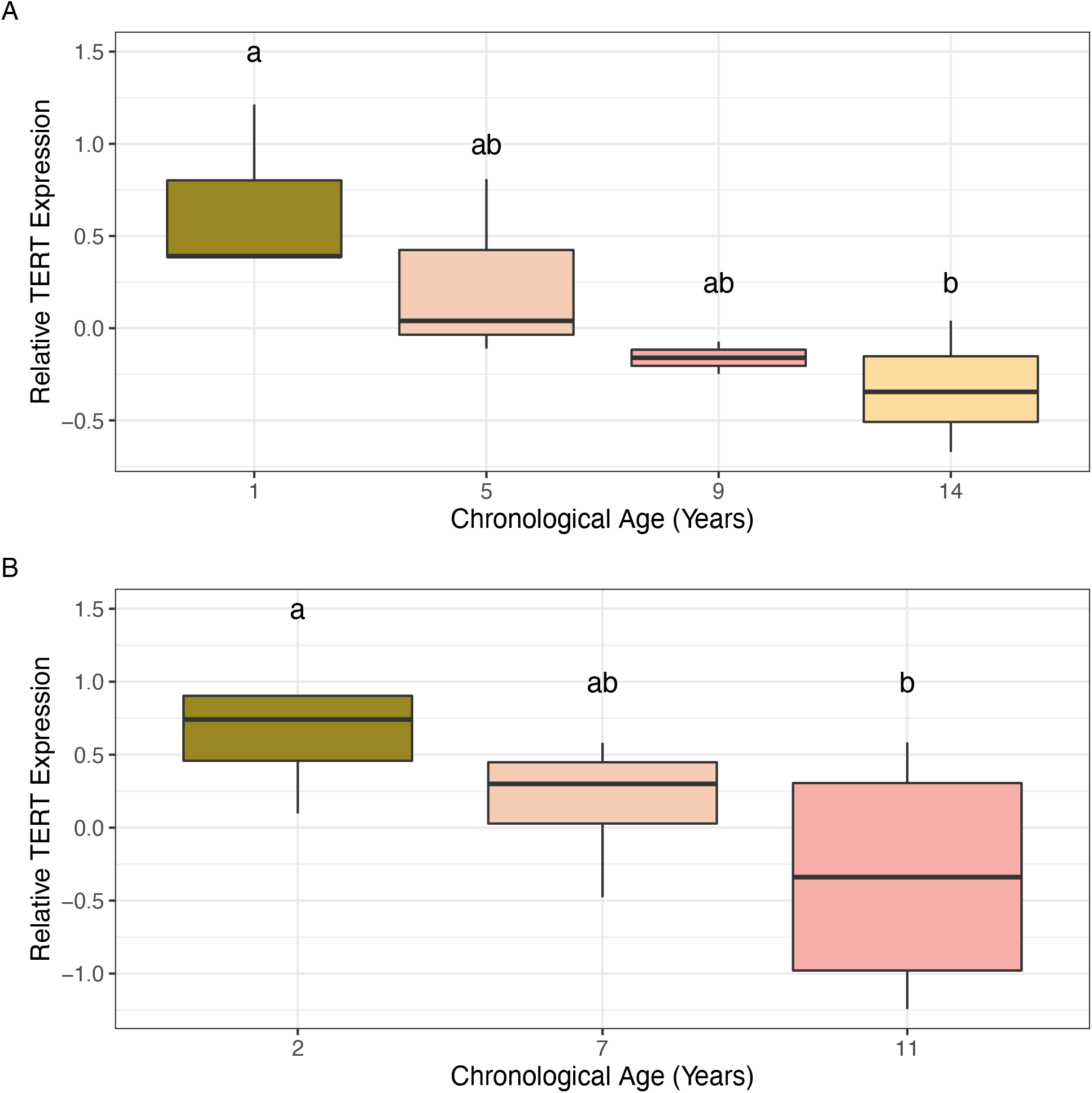
Normalized expression of *TERT* for almond samples within the age cohorts test. **(a)** Age cohort collected in 2018. **(b)** Age cohort collected in 2019. Significant differences in relative expression between age cohorts based on ANOVA followed by *post hoc* Tukey’s HSD (alpha = 0.1) are denoted by the letter groupings (ANOVA 2018 p-value = 0.09087; ANOVA 2019 p-value = 0.1414).

## Discussion

Almond, an economically-valuable nut crop, exhibits an aging-related disorder known as non-infectious bud failure that negatively impacts vegetative development and ultimately, yield. As a clonally propagated crop, tracking age and thus susceptibility to bud failure is difficult, making biomarkers of age a valuable resource to circumvent the impacts of aging-related disorders in almond germplasm. Telomere length is used as a biomarker of age and development of age-related disorders in mammals, but the association between telomere length and age in plants is not well-defined (Watson and Riha, 2011; Procházková Schrumpfová *et al*., 2019). The present study tests the hypothesis that telomere length and/or *TERT* expression are associated with age in almond. To test this, both *in silico* and qPCR approaches were utilized to measure telomere length and estimate *TERT* expression in sets of almond accessions of known chronological age. *In silico* analysis was performed using whole-genome sequencing data from nine almond accessions of known age. Samples were collected from three and four sets of age cohorts over two years to test for an association between relative telomere length and individual age using the MMQPCR method as well as between *TERT* expression and age using qRT-PCR.

### In silico and quantitative PCR approaches suggest a marginal association between average or relative telomere length and age in almond

Average telomere length estimated *in silico* using the program Computel (Nersisyan and Arakelyan, 2015) revealed a non-significant, negative association with age in the almond accessions tested. This same pattern was shown utilizing MMQPCR and almond leaf samples collected from different almond age cohorts in 2018 and 2019 where telomere length decreases with increasing age. The association demonstrated in this study adds to the growing body of knowledge regarding the complex relationship between telomere length and plant aging.

Previous studies in both *Ginkgo biloba* and *Panax ginseng* showed a pattern of increased telomere length with increased age, suggesting plants do not follow the same patterns of telomere shortening as seen in mammals (Liang *et al*., 2015; Liu *et al*., 2007). Work in apple (*Malus domestica*) and *Prunus yedoensis*, both members of Rosaceae like almond, show no change in telomere lengths with increased plant age over a five year timespan (Moriguchi *et al*., 2007). In bristlecone pine (*Pinus longaeva*), a long-lived perennial gymnosperm, telomere lengths measured in needle and root tissues between 0 – 3,500 years old showed a cyclical pattern of lengthening and shortening with age (Flanary and Kletetschka, 2005). Further, when analyzing telomere length in relation to tissue differentiation, studies in both barley (*Hordeum vulgare*) and Scots pine (*Pinus sylvestris*) showed telomere shortening through embryo development to leaf or needle formation (Aronen and Ryynänen, 2012; Kilian *et al*., 1995). Similarly, in silver birch (*Betula pendula*), telomeres shorten when plant are grown in tissue culture conditions compared to those grown outdoors, suggesting abiotic stressors may induce telomere shortening (Aronen and Ryynänen, 2014).

The results in almond suggest a pattern closest to what was observed in bristlecone pine where telomere lengths shorten and lengthen throughout an individual’s lifetime. This pattern could be unique to gymnosperms, however, and needs to be further characterized in angiosperms including Rosaceous species. While the commercial lifespan of almond is typically less than 30 years, almond can live more than 150 years (Micke, 1996). In this study, the maximum age tested via the *in silico* approach was 32 years-old and via qPCR was 14 years-old, suggesting that a wider age-range of trees and a larger sample size could produce a more refined model of telomere length patterns over time.

Current almond cultivars may also be ontogenetically old, such as ‘Nonpareil’, the most relevant US cultivar representing ∼40% of acreage, which was first described almost 140 years ago and has been propagated by budding since (Wickson, 1914; Almond Board of California, 2019). The ontogenetic age of a cultivar may be a factor to consider in the onset of aging-related disorders like BF in almond. Additionally, it would be interesting to track the change in telomere length following clonal propagation (through budding) in which plants experience a rejuvenation process, reverting to a juvenile state for a short period of time (Bonga, 1982). It was further found that propagating almond from basal epicormic buds, potentially representing ontogenetically young meristematic tissue, seemed to alleviate BF in resulting clones (Gradziel *et al*., 2019). Testing telomere lengths in epicormic tissues could present another avenue to both track aging in almond and develop biomarkers to predict BF potential in almond.

### TERT expression measured by qRT-PCR is a putatively associated with age in almond accessions

To test the hypothesis that *TERT* expression can serve as a biomarker of age in almond, expression patterns were tested in cohorts representing either three or four distinct ages over two years. Results from this work showed a consistent pattern of marginally significant, decreased expression with increased ontogenetic age. Telomerase was shown to be a modulator of longevity in humans and other mammals, but work describing telomerase patterns in plants is limited (Boccardi and Paolisso, 2014; Fitzgerald *et al*., 1996).

A comprehensive study examining telomerase protein activity in carrot (*Daucus carota*), cauliflower (*Brassica oleracea*), soybean (*Glycine max*), *Arabidopsis thaliana*, and rice (*Oryza sativa*) demonstrated that, like telomere lengths, protein activity tends to be highest in undifferentiated tissues like meristematic tissues and is lower in differentiated tissues such as leaves (Fitzgerald *et al*., 1996). This result was supported by further work in barley and maize showing little activity in differentiated tissues (Kilian *et al*., 1998). These studies were all performed in annuals or biennials, however, suggesting that telomerase activity does in fact decrease with increased plant age in these crops. Work in perennials including bristlecone pine, *P. ginseng*, and *G. biloba* showed an association between telomerase activity and age, suggesting patterns unique to perennial plant species (Liang *et al*., 2015; Flanary and Kletetschka, 2005; Song *et al*., 2011). A study in almond could be performed using a wider age-range and larger sample size to elucidate the effect of age on telomerase activity, similar to what was referenced above for telomere length measurements. Additionally, many of the studies performed in other plants examining patterns of telomerase activity focused on protein activity rather than gene expression. A future study will be necessary in almond to examine the telomerase protein activity potentially by Western blot or other proteomics approaches to corroborate the association between *TERT* expression and protein activity.

While a pattern was established in plants demonstrating a direct relationship between telomerase activity and telomere length, regulation of telomerase is still not well understood in the plant kingdom (Zachová *et al*., 2013; Jurečková *et al*., 2017; Fitzgerald *et al*., 1996). Interestingly, work in Arabidopsis has shown a link between DNA methylation and telomere length, suggesting that this epigenetic mark likely has a role in regulating telomere lengths potentially by modulating telomerase activity (Vega-Vaquero *et al*., 2016; Lee and Cho, 2019; Ogrocká *et al*., 2014). A study is ongoing in almond to analyze DNA methylation patterns in a set of almond accessions representing three distinct age cohorts to determine what, if any, impact age has on methylation profiles.

Despite the limited age-range and small sample size used in this study, a consistent pattern of both decreased telomere length and decreased *TERT* expression with increased age was observed over two years of sampling. These results provide a basis for future study and exploration into the utility of telomere length measurement and/or *TERT* expression or telomerase activity as biomarkers of aging in almond. Developing a robust biomarker to track aging in almond, a primarily clonally propagated crop, would allow growers, producers, and breeders to screen germplasm to eliminate selections or clones with a high susceptibility to age-related disorders due to advanced ontogenetic age.

## Supplementary Files

**S1**. File containing raw data for each experiment including *in silico* telomere length estimation, mean telomere length quantification, and *TERT* expression analysis.

**S2**. R code used to perform all statistical analyses reported in this manuscript.

## Acknowledgements

We would like to acknowledge Matthew Willman for his assistance with the statistical analyses and Cheri Nemes for her assistance with wet lab portions of this project. We would like to thank Daniel Williams for editing later versions of this manuscript. This work is supported by the Ohio State University CFAES-SEEDS program 2018113, the Translational Plant Sciences Graduate Fellowship, the AFRI-EWD Predoctoral Fellowship 2019-67011-29558 from the USDA National Institute of Food and Agriculture, and the Ohio Supercomputer Center.

## References

Alioto, T., Alexiou, K.G., Bardil, A., et al. (2020) Transposons played a major role in the diversification between the closely related almond and peach genomes: results from the almond genome sequence. The Plant Journal, 101, 455–472. 10.1111/tpj.14538.

Almond Board of California (2019) Almond Almanac, Modesto, CA: Almond Board of California.

Anchelin, M., Murcia, L., Alcaraz-Pérez, F., García-Navarro, E.M. and Cayuela, M.L. (2011) Behaviour of telomere and telomerase during aging and regeneration in zebrafish. PLoS One, 6. 10.1371/journal.pone.0016955.

Aronen, T. and Ryynänen, L. (2014) Silver birch telomeres shorten in tissue culture. Tree Genetics & Genomes, 10, 67–74. 10.1007/s11295-013-0662-4.

Aronen, T. and Ryynänen, L. (2012) Variation in telomeric repeats of Scots pine (*Pinus sylvestris L.*). Tree Genetics & Genomes, 8, 267–275. 10.1186%2F1753-6561-5-S7-O42.

Aviv, A. and Shay, J.W. (2018) Reflections on telomere dynamics and ageing-related diseases in humans. Philos. Trans. R. Soc. Lond., B, Biol. Sci., 373. 10.1098/rstb.2016.0436.

Bastias, A., Oviedo, K., Almada, R., Correa, F. and Sagredo, B. (2020) Identifying and validating housekeeping hybrid *Prunus* spp. genes for root gene-expression studies. PLOS ONE, 15. 10.1371/journal.pone.0228403.

Boccardi, V. and Paolisso, G. (2014) Telomerase activation: A potential key modulator for human healthspan and longevity. Ageing Research Reviews, 15, 1–5. 10.1016/j.arr.2013.12.006.

Bonga, J.M. (1982) Vegetative propagation in relation to juvenility, maturity, and rejuvenation. In J. M. Bonga and D. J. Durzan, eds. Tissue Culture in Forestry. Forestry Sciences. Dordrecht: Springer Netherlands, pp. 387–412. 10.1007/978-94-017-3538-4_13.

Brutovská, E., Sámelová, A., Dušička, J. and Mičieta, K. (2013) Ageing of trees: Application of general ageing theories. Ageing Research Reviews, 12, 855–866. 10.1016/j.arr.2013.07.001.

Bustin, S.A., Benes, V., Garson, J.A., et al. (2009) The MIQE guidelines: minimum information for publication of quantitative real-time PCR experiments. Clin. Chem., 55, 611–622. 10.1373/clinchem.2008.112797.

Cawthon, R.M. (2009) Telomere length measurement by a novel monochrome multiplex quantitative PCR method. Nucleic Acids Res, 37, e21. 10.1093/nar/gkn1027.

De la Torre-Espinosa, Z.Y., Barredo-Pool, F., Castaño de la Serna, E. and Sánchez-Teyer, L.F. (2020) Active telomerase during leaf growth and increase of age in plants from *Agave tequilana* var. Azul. Physiology and Molecular Biology of Plants, 26, 639-647. 10.1007/s12298-020-00781-7.

Fitzgerald, M.S., McKnight, T.D. and Shippen, D.E. (1996) Characterization and developmental patterns of telomerase expression in plants. PNAS, 93, 14422–14427. 10.1073/pnas.93.25.14422.

Fitzgerald, M.S., Riha, K., Gao, F., Ren, S., McKnight, T.D. and Shippen, D.E. (1999) Disruption of the telomerase catalytic subunit gene from *Arabidopsis* inactivates telomerase and leads to a slow loss of telomeric DNA. PNAS, 96, 14813–14818. 10.1073/pnas.96.26.14813.

Flanary, B.E. and Kletetschka, G. (2005) Analysis of telomere length and telomerase activity in tree species of various life-spans, and with age in the bristlecone pine *Pinus longaeva*. Biogerontology, 6, 101–111. 10.1007/s10522-005-3484-4.

Fossel, M. (1998) Telomerase and the aging cell: Implications for human health. JAMA, 279, 1732–1735. 10.1001/jama.279.21.1732.

Gambino, G., Perrone, I. and Gribaudo, I. (2008) A Rapid and effective method for RNA extraction from different tissues of grapevine and other woody plants. Phytochemical Analysis, 19, 520–525. 10.1002/pca.1078.

Gradziel, T., Lampinen, B. and Preece, J.E. (2019) Propagation from basal epicormic meristems remediates an aging-related disorder in almond clones. Horticulturae, 5, 28. 10.3390/horticulturae5020028.

Gradziel, T.M., Thorpe, M.A., Fresnedo-Ramírez, J., et al. (2013) Molecular marker based diagnostics for almond non-infectious bud-failure, Almond Board of California.

Hemann, M.T., Strong, M.A., Hao, L.-Y. and Greider, C.W. (2001) The shortest telomere, not average telomere length, is critical for cell viability and chromosome stability. Cell, 107, 67–77. 10.1016/s0092-8674(01)00504-9.

Initiative, T.I.P.G., Verde, I., Abbott, A.G., et al. (2013) The high-quality draft genome of peach (*Prunus persica*) identifies unique patterns of genetic diversity, domestication and genome evolution. Nature Genetics, 45, 487–494. 10.1038/ng.2586.

Jurečková, J.F., Sýkorová, E., Hafidh, S., Honys, D., Fajkus, J. and Fojtová, M. (2017) Tissue-specific expression of telomerase reverse transcriptase gene variants in Nicotiana tabacum. Planta, 245, 549–561. 10.1007/s00425-016-2624-1.

Kester, D.E. and Jones, R.W. (1970) Non-infectious bud-failure from breeding programs of almond (*Prunus amygdalus* Batsch). Journal of the American Society of Horticultural Science, 95, 492–96.

Kester, D.E., Shackel, K.A., Micke, W.C., Viveros, M. and Gradziel, T.M. (2004) Noninfectious bud failure in ‘Carmel’ almond: I. Pattern of development in vegetative progeny trees. J. Amer. Soc. Hort. Sci., 129, 244–249. 10.21273/JASHS.129.2.0244.

Kilian, A., Heller, K. and Kleinhofs, A. (1998) Development patterns of telomerase activity in barley and maize. Plant Mol Biol, 37, 621–628. 10.1023/A:1005994629814.

Kilian, A., Stiff, C. and Kleinhofs, A. (1995) Barley telomeres shorten during differentiation but grow in callus culture. Proc Natl Acad Sci USA, 92, 9555. 10.1073/pnas.92.21.9555.

Kirkwood, T.B.L. (2005) Understanding the odd science of aging. Cell, 120, 437–447. 10.1016/j.cell.2005.01.027.

Lee, W.K. and Cho, M.H. (2019) Epigenetic aspects of telomeric chromatin in *Arabidopsis thaliana*. BMB Rep, 52, 175–180. 10.5483/BMBRep.2019.52.3.047.

Liang, J., Jiang, C., Peng, H., Shi, Q., Guo, X., Yuan, Y. and Huang, L. (2015) Analysis of the age of *Panax ginseng* based on telomere length and telomerase activity. Sci Rep, 5. 10.1038/srep07985.

Liu, D., Qiao, N., Song, H., Hua, X., Du, J., Lu, H. and Li, F. (2007) Comparative analysis of telomeric restriction fragment lengths in different tissues of *Ginkgo biloba* trees of different age. Journal Of Plant Research, 120, 523–528. 10.1007/s10265-007-0092-1.

Marioni, R.E., Harris, S.E., Shah, S., McRae, A.F., Zglinicki, T. von, Martin-Ruiz, C., Wray, N.R., Visscher, P.M. and Deary, I.J. (2016) The epigenetic clock and telomere length are independently associated with chronological age and mortality. Int J Epidemiol, 45, 424–432. 10.1093/ije/dyw041.

Micke, W.C. (1996) Almond Production Manual, UCANR Publications.

Montpetit, A.J., Alhareeri, A.A., Montpetit, M., et al. (2014) Telomere length: A review of methods for measurement. Nurs Res, 63, 289–299. 10.1097/NNR.0000000000000037.

Moriguchi, R., Kato, K., Kanahama, K., Kanayama, Y. and Kikuchi, H. (2007) Analysis of telomere lengths in apple and cherry trees. Acta Horticulturae, 389–395. 10.17660/ActaHortic.2007.738.47.

Munné-Bosch, S. (2007) Aging in perennials. Critical Reviews in Plant Sciences, 26, 123–138. 10.1080/07352680701402487.

Nelson, A.D.L., Beilstein, M.A. and Shippen, D.E. (2014) Plant telomeres and telomerase. In S. H. Howell, ed. Molecular Biology. New York, NY: Springer New York, pp. 25–49. 10.1007/978-1-4614-7570-5_4.

Nersisyan, L. and Arakelyan, A. (2015) Computel: Computation of mean telomere length from whole-genome next-generation sequencing data. PLOS ONE, 10. 10.1371/journal.pone.0125201.

Ogrocká, A., Polanská, P., Majerová, E., Janeba, Z., Fajkus, J. and Fojtová, M. (2014) Compromised telomere maintenance in hypomethylated *Arabidopsis thaliana* plants. Nucleic Acids Res, 42, 2919–2931. 10.1093/nar/gkt1285.

Oguchi, K., Liu, H., Tamura, K. and Takahashi, H. (1999) Molecular cloning and characterization of AtTERT, a telomerase reverse transcriptase homolog in *Arabidopsis thaliana*. FEBS Letters, 457, 465–469. 10.1016/s0014-5793(99)01083-2.

Ohio Supercomputer Center. 1987. Ohio Supercomputer Center. Columbus OH: Ohio Supercomputer Center. http://osc.edu/ark:/19495/f5s1ph73.

Poethig, R.S. (2003) Phase change and the regulation of developmental timing in plants. Science, 301, 334–336. 10.1126/science.1085328.

Procházková Schrumpfová, P., Fojtová, M. and Fajkus, J. (2019) Telomeres in plants and humans: Not so different, not so similar. Cells, 8, 58. 10.3390/cells8010058.

Runov, A.L., Vonsky, M.S. and Mikhelson, V.M. (2015) DNA methylation level and telomere length as a basis for modeling of the biological aging clock. Cell and Tissue Biology, 9, 261–264. 10.1134/S1990519X15040094.

Sanders, J.L. and Newman, A.B. (2013) Telomere length in epidemiology: A biomarker of aging, age-related disease, both, or neither? Epidemiol Rev, 35, 112–131. 10.1093/epirev/mxs008.

Song, H., Liu, D., Li, F. and Lu, H. (2011) Season-and age-associated telomerase activity in *Ginkgo biloba* L. Mol Biol Rep, 38, 1799–1805. 10.1007/s11033-010-0295-8.

Sweetlove, L. and Gutierrez, C. (2019) The journey to the end of the chromosome: delivering active telomerase to telomeres in plants. Plant J, 98, 193–194. 10.1111/tpj.14328.

Thomas, H. (2013) Senescence, ageing and death of the whole plant. New Phytol., 197, 696–711. 10.1111/nph.12047.

Tong, Z., Gao, Z., Wang, F., Zhou, J. and Zhang, Z. (2009) Selection of reliable reference genes for gene expression studies in peach using real-time PCR. BMC Mol Biol, 10, 71. 10.1186/1471-2199-10-71.

Vaquero-Sedas, M.I. and Vega-Palas, M.A. (2014) Determination of *Arabidopsis thaliana* telomere length by PCR. Scientific Reports, 4. 10.1038/srep05540.

Vega-Vaquero, A., Bonora, G., Morselli, M., Vaquero-Sedas, M.I., Rubbi, L., Pellegrini, M. and Vega-Palas, M.A. (2016) Novel features of telomere biology revealed by the absence of telomeric DNA methylation. Genome Res, 26, 1047–1056. 10.1101/gr.202465.115.

Verhulst, S. (2020) Improving comparability between qPCR-based telomere studies. Molecular Ecology Resources, 20, 11–13. 10.1111/1755-0998.13114.

Wang, T., Hao, R., Pan, H., Cheng, T. and Zhang, Q. (2014) Selection of suitable reference genes for quantitative real-time polymerase chain reaction in *Prunus mume* during flowering stages and under different abiotic stress conditions. J. Amer. Soc. Hort. Sci., 139, 113–122. 10.21273/JASHS.139.2.113.

Watson, J.M. and Riha, K. (2011) Telomeres, aging, and plants: From weeds to Methuselah – a mini-review. GER, 57, 129–136. 10.1159/000310174.

Wickson, E.J. (1914) The California fruits and how to grow them, San Francisco, CA: Pacific rural Press. https://lccn.loc.gov/14011514.

Zachová, D., Fojtová, M., Dvořáčková, M., Mozgová, I., Lermontova, I., Peška, V., Schubert, I., Fajkus, J. and Sýkorová, E. (2013) Structure-function relationships during transgenic telomerase expression in *Arabidopsis*. Physiol Plantarum, 149, 114–126. 10.1111/ppl.12021.

Zangi, M., Bagherieh Najjar, M.B., Golalipour, M. and Aghdasi, M. (2019) met1 DNA methyltransferase controls TERT gene expression: A new insight to the role of telomerase in development. CellJ, 22. 10.22074/cellj.2020.6290.

