## Supplementary File 2 for "Telomere length and *TERT* expression are associated with age in almond (*Prunus dulcis* [Mill.] D.A.Webb)"

#In silico mean telomere length estimation linear regression analysis
#Load required packages
require(ggplot2)
require(wesanderson)
require(ggrepel)
require(ggpubr)

#Read in output from Computel in file 'ComputelOutput.csv'; file headers: Age (years), meanTelo (mean telomere length - basepairs), Name (cultivar or identifier)

telomereData <- read.csv('ComputelOutput.csv')

#Perform square root transformation on mean telomere length measurements

telomereData$sqrt <- sqrt(telomereData$meanTelo)

#Perform Shapiro-Wilk normality test on square root-transformed mean telomere length to assess normality assumptions

shapiro.test(telomereData$sqrt)

#Regress square root-transformed mean telomere length on almond age

linearMod <- lm(sqrt ~ Age, data=telomereData)

#Display R^2 and p-value for the linear model

summary(linearMod2) 

#Create regression plot displaying almond accession Names and best-fit line

ggplot(telomereData, aes(x = Age, y = sqrt)) +
  geom_point(size = 2.5) +
  stat_smooth(method = "lm",
              col = wes_palette("Royal2", 1),
              se = FALSE,
              size = 1, linetype = "dashed") +
  theme_bw() +
  labs(x = "Chronological Age (Years)", y = "SQRT Mean Telomere Length (Base Pairs)" ) +
  geom_label_repel(aes(label = Name),
                   box.padding   = 0.35, 
                   point.padding = 0.5,
                   segment.color = 'grey50') + stat_regline_equation(label.x = 23, label.y = 31, aes(size = 16, fontface = "bold.italic"), show.legend = FALSE)

#Analysis of mean telomere lengths generated by monochrome multiplex qPCR comparing distinct almond age cohorts collected in 2018 and 2019
#Load required packages
require(agricolae)
require(ggplot2)
require(wesanderson)
require(multipanelfigure)
require(grid)

#Read in mean telomere length data from file 'TelomereData.csv'; file headers: Year (year planted), Replicate (biological replicate number), T.S (estimated T/S ratio genearted from standard curve using telomere and RPII Cq values)

telo_data <- read.csv("TelomereData.csv")

#Convert variable 'Replicate' to a factor for analysis

telo_data$Replicate <- as.factor(telo_data$Replicate)

#Calculate the average T/S ratio for the reference almond sample to use to calculate relative telomere lengths for each almond indivdiual

referenceAverage <- mean(c(0.4758955, 0.8377413, 0.779102))

#Create a relative telomere length variable (rTL)

telo_data$rTL <- telo_data$T.S/referenceAverage

#Create a z-score variable (zscore_rTL) by calculating the z-score for each almond sample

telo_data$zscore_rTL <- (telo_data$rTL - mean(telo_data$rTL)/sd(telo_data$rTL))

#Create a variable called age listed the chronological age of each almond in years; make this variable a factor
#Example is for data collected in 2019

telo_data$age <- c(2,2,2,7,7,7,11,11,11)

telo_data$age <- as.factor(telo_data$age)

#Perform Shapiro-Wilk normality test on zscore_rTL assess test normality assumption

shapiro.test(telo_data$zscore_rTL)

#Perform Bartlett test to assess homogeneity of variance assumption

bartlett.test(zscore_rTL ~ age, data = telo_data)

#Regress zscore_rTL on almond age

fit <- lm(telo_data$zscore_rTL ~ telo_data$age)

#Perform ANOVA to generate p-value on linear model

anova(fit)

#Perform post-hoc Fisher's LSD for means separation

LSD_output <- LSD.test(fit6, "telo_data$age", alpha = 0.1, p.adj = "none")
LSD_output

#Create coords variable to display letter groupings based on means separation results
#Example is for data collected in 2019

coords <- data.frame("x" = c(1,2,3), "y" = c(1.5,0,-0.5), "sig" = c("a", "ab", "b"))

#Create boxplot displaying the zscore of relative telomere lenght (zscore_rTL) for each almond age cohort, including the means separation groupings
#Example is for data collected in 2019

Telo19 <- ggplot(telo_data, aes(x=age, y=zscore_rTL, fill = age)) + geom_boxplot() + 
  labs(x = "Chronological Age (Years)", y = "Zscore Relative Telomere Length") +
  theme_bw()+
  theme(legend.position = "none", text = element_text(size = 12)) +
  geom_text(data = coords, mapping = aes(x=x, y=y, label = sig, size = 12), inherit.aes = FALSE) +
  scale_fill_manual(values=wes_palette(name="Royal2"))

#Create a multipanel figure (figure1) combining boxplot produced with 2018 data and boxplot produced with 2019 data
figure1 <- multi_panel_figure(width = 180, height = 180, columns = 1, rows = 2)
figure1 %<>% fill_panel(Telo18)
figure1 %<>% fill_panel(Telo19)
figure1

#Analysis of relative TERT expression data generated via qRT-PCR comparing distinct almond age cohorts collected in 2018 and 2019
#Load required packages
require(ggplot2)
require(agricolae)
require(multipanelfigure)
require(grid)

#Read in relative expression data from file 'TERTExpression.csv'; file headers: Chronological.Age (age of each almond accession in years), Expression (normalized relative TERT expression)

TERT_data <- read.csv('TERTExpression.csv')

#Convert variable 'Chronological.Age' to a factor

TERT_data$Chronological.Age <- as.factor(TERT_data$Chronological.Age)

#Perform a log2 transformation on 'Expression' data variable

TERT_data$logExpression <- log(TERT_data$Expression)

#Omit any missing data values (n/a's)

TERT_data <- na.omit(TERT_data)

#Perform Shapiro-Wilk normality test on logExpression to assess test normality assumption

shapiro.test(TERT_data$logExpression)

#Perform Bartlett test to assess homogeneity of variance assumption

bartlett.test(logExpression ~ Chronological.Age, data = TERT_data)

#Regress log-transformed Expression on Chronological.Age

fit_tert <- lm(log(TERT_data$Expression) ~ TERT_data$Chronological.Age)

#Perform ANOVA to generate p-value on linear model

anova(fit_tert)

#Perform post-hoc Tukey's HSD for means separation

HSD_tert <- HSD.test(fit_tert, "TERT_data$Chronological.Age", alpha = 0.1)
HSD_tert

#Create coords variable to display letter groupings based on means separation results
#Example is for data collected in 2018

tert_coords <- data.frame("x" = c(1,2,3,4), "y" = c(1.5,1,0.25,0.25), "sig" = c("a", "ab", "ab", "b"))

#Create boxplot displaying the zscore of relative telomere lenght (zscore_rTL) for each almond age cohort, including the means separation groupings
#Example is for data collected in 2018

TERT2018 <- ggplot(TERT_data, aes(x=Chronological.Age, y=logExpression, fill = Chronological.Age)) + 
  geom_boxplot() +
  labs(x = "Chronological Age (Years)", y = "Relative TERT Expression") +
  theme_bw()+
  theme(legend.position = "none", text = element_text(size = 12)) +
  geom_text(data = tert_coords, mapping = aes(x=x, y=y, label = sig, size = 12), inherit.aes = FALSE) +
  scale_fill_manual(values=wes_palette(name="Royal2"))

#Create a multipanel figure (figure2) combining boxplot produced with 2018 data and boxplot produced with 2019 data

figure2 <- multi_panel_figure(width = 180, height = 180, columns = 1, rows = 2)
figure2 %<>% fill_panel(TERT2018)
figure2 %<>% fill_panel(TERT2019)
figure2
